# Targeted photodynamic neutralization of SARS-CoV-2 mediated by singlet oxygen

**DOI:** 10.1101/2022.11.29.518438

**Authors:** Ruhui Yao, Jian Hou, Xin Zhang, Yi Li, Junhui Lai, Qinqin Wu, Qinglian Liu, Lei Zhou

## Abstract

The SARS-CoV-2 virus has been on a rampage for more than two years. Vaccines in combination with neutralizing antibodies (NAbs) against SARS-CoV-2 carry great hope in the treatment and final elimination of COVID-19. However, the relentless emergence of variants of concern (VOC), including the most recent Omicron variants, presses for novel measures to counter these variants that often show immune evasion. Hereby we developed a targeted photodynamic approach to neutralize SARS-CoV-2 by engineering a genetically encoded photosensitizer (SOPP3) to a diverse list of antibodies targeting the WT spike protein, including human antibodies isolated from a 2003 SARS patient, potent monomeric and multimeric nanobodies targeting RBD, and non-neutralizing antibodies (non-NAbs) targeting the more conserved NTD region. As confirmed by pseudovirus neutralization assay, this targeted photodynamic approach significantly increased the efficacy of these antibodies, especially that of non-NAbs, against not only the WT but also the Delta strain and the heavily immune escape Omicron strain (BA.1). Subsequent measurement of infrared phosphorescence at 1270 nm confirmed the generation of singlet oxygen (^1^O_2_) in the photodynamic process. Mass spectroscopy assay uncovered amino acids in the spike protein targeted by ^1^O_2_. Impressively, Y145 and H146 form an oxidization “hotspot”, which overlaps with the antigenic “supersite” in NTD. Taken together, our study established a targeted photodynamic approach against the SARS-CoV-2 virus and provided mechanistic insights into the photodynamic modification of protein molecules mediated by ^1^O_2_.

## INTRODUCTION

Photodynamic therapy (PDT) is an FDA-approved procedure to treat several types of cancer and other conditions including infections caused by pathogens.^1–3^ Three elements are involved in PDT: photosensitizer, oxygen, and light. Photosensitizers in the ground state receive energy from the excitation light to produce singlet oxygen (^1^O_2_) through type II photosensitized process or other ROS through the type I process.^4^ ^1^O_2_ is the molecular oxygen in electronically excited states and is chemically very reactive, and directly oxidizes molecules in the cell including protein, DNA, and lipid. Excessive ^1^O_2_ is toxic and causes damage to biomolecules and cellular structures, which has been utilized in basic research as the Chromophore-Assisted Light Inactivation (CALI) approach and in clinical practice as PDT.^5, 6^ Due to the unique physicochemical properties of ^1^O_2_, namely its brief lifetime in microseconds and short working distance in nanometers, CALI and PDT can achieve precise elimination of target molecules or cells without any collateral damage. Significant benefits of PDT include fewer side effects, much reduced long-term morbidity, high effectiveness, increased selectivity, lower cost, and minimal invasiveness. Notably, site-specific delivery of photosensitizers to the target cell or tissue is the critical factor in PDT.

As an extension and branch of PDT, photodynamic antimicrobial chemotherapy (PACT) also utilizes the ^1^O_2_ of photosensitized production and can effectively eliminate pathogens including viruses such as HPV and HIV, fungi, and bacteria.^3, 7, 8^ Compared to human cells, these microorganisms are generally of smaller size and lack complicated intracellular membrane structures which form layers of extra protection, therefore they are more vulnerable to photochemical toxicity.^9, 10^ In recent years, PACT has received much attention in treating drug-resistant germs, which represents an ever-increasing threat to public health due to the widespread usage of antibiotics.^3, 7^ Just like PDT, “smart” photosensitizers that specifically recognize the target cell critically determine the effectiveness of PACT.^9, 11^

Caused by the SARS-CoV-2 virus, the coronavirus disease 2019 (COVID-19) was first reported in late 2019 and has become a pandemic for more than two years. The SARS-CoV-2 virus belongs to the family of positive-sense single-stranded RNA viruses, which also include seasonal coronavirus, SARS-CoV-1 (SARS-CoV), and Middle East Respiratory Syndrome (MERS)-CoV.^12^ SARS-CoV-2 attacks cells in the respiratory system, mainly through a specific interaction between the spike glycoprotein on the viral surface and the corresponding receptor on the cell surface, the angiotensin-converting enzyme 2 (ACE2).^13, 14^ Therefore, much of the ongoing research efforts against COVID-19 have been devoted to understanding the spike protein in membrane fusion and viral entry and to developing strategies to block the interaction between spike with ACE2.^15, 16^ The spike protein of SARS-CoV-2 is composed of three presumably identical protomers, with each protomer containing 1273 amino acids. The full-length S protein includes an extracellular ectodomain, a transmembrane domain (a.a. 1213-1237), and a short cytoplasmic tail. The mature form of the spike protein is extensively glycosylated, with 22 potential N-glycosites and numerous O-glycosites decorating the ectodomain. A distinctive polybasic protease (furin) cleavage site separates the S protein into S1 (a.a. 1-685) and S2 (a.a. 686-1273).^17^ In addition, within the S2 domain, the S2’ site can be cleaved by TMPRSS2, a membrane-anchoring serine protease.^18^ These two proteolytic processes play important roles in the life cycle of SARS-CoV-2. It is believed that the S1 contributes to the initial contact with ACE2 on the host cell surface, whereas S2 is responsible for the following membrane fusion and viral entry into the host cell. Within the S1 domain, two structural motifs stand out: the more conserved N-terminal domain (NTD, a.a. 13-305) and the receptor-binding domain (RBD, a.a. 319-541). RBD is in direct contact with ACE2 and therefore play a dominant role in the initial viral recognition of host cell. Driven by a concerted effort by the research community, our understanding of RBD and its interaction with ACE2 has been advancing rapidly. RBD adopts two dramatic conformations, down (closed) and up (open) states.^19^ In the open state, RBD swings upward and interacts with the ACE2 through a patch of ~25 amino acids.

Antibodies that target the spike protein, especially RBD, can effectively block the interaction between spike and ACE2 and thereby neutralize the virus from infecting host cells. During the past two years, neutralizing antibodies (NAbs) against SARS-CoV-2 have been identified from multiple sources, including convalescent patients recovering from infections of SARS-CoV-2, saved blood cells collected from individuals infected with SARS-CoV-1 in 2003, naïve human antibody libraries, and animals including alpacas and genetically humanized mice (VelocImmune) infected with SARS-CoV-2. As of February 1^st^, 2022, the database of CoV-AbDab has collected over 5000 antibodies (including nanobodies) against SARS-CoV, SARS-CoV-2, and MERS-CoV.^20^ However, mutant SARS-CoV-2 strains, especially the most recent Omicron variant that carries extensive mutations in RBD, evade the majority of existing NAbs developed so far.^21–23^ Tremendous pressure has been landed on the research community and industry to update the current treatment strategy against SARS-CoV-2.^24–26^ In contrast to NAbs, non-neutralizing antibodies (non-NAbs) do not produce much selective pressure for SARS-CoV-2, and among them, the antibodies that target conserved regions in spike have a broad recognition for diverse variants. The application potential of these non-NAbs still requires further exploration.

Recently published studies have explored the effectiveness and potential of PDT in the neutralization of SARS-CoV-2.^27^ However, those studies used non-specific photosensitizers, and long exposure of intense excitation light was required.^28, 29^ We set out to develop a targeted photodynamic approach to neutralize the SARS-CoV-2 virus. We utilized both NAbs and non-NAbs reported in the literature during the past two years and tagged them with a genetically encoded photosensitizer. Our results showed that brief light pulses effectively increased neutralization efficacy against not only the wild-type but also the Delta and Omicron strains. Moreover, among the oxidatively modified residues identified by mass-spectrometry, two significant residues are located in the center of the antigenic “supersite” discovered in the NTD domain.

## RESULTS

To explore the applicability of targeted photodynamic neutralization against SARS-CoV-2, we chose antibodies that target either RBD or NTD of the WT spike protein and with different functionality. We examined not only NAbs but also non-neutralizing and even infection enhancing antibodies: 1) S309 (potent and broad-spectrum) and S304, two neutralizing human antibodies targeting RBD, isolated from a 2003 SARS-CoV patient^30^ (**Fig. 1a**); 2), mNb6 (a matured form with a 500-fold increase in binding affinity) and mNb6-tri (a trivalent form with no detectable dissociation from the spike protein), two neutralizing nanobodies targeting RBD, isolated from a screening of a yeast surface-displayed library^31^; 3) DH1052, DH1054, DH1055, and DH1056, four non-neutralizing human antibodies targeting NTD, isolated from a SARS-CoV-2 patient^32^; 4) WNb58, a non-neutralizing nanobody recognizing the RBDs of both SARS-CoV-1 and SARS-CoV-2, isolated from two alpacas immunized with SARS-CoV-1 RBD, SARS-CoV-2 spike and RBD^33^.

**Fig. 1.**
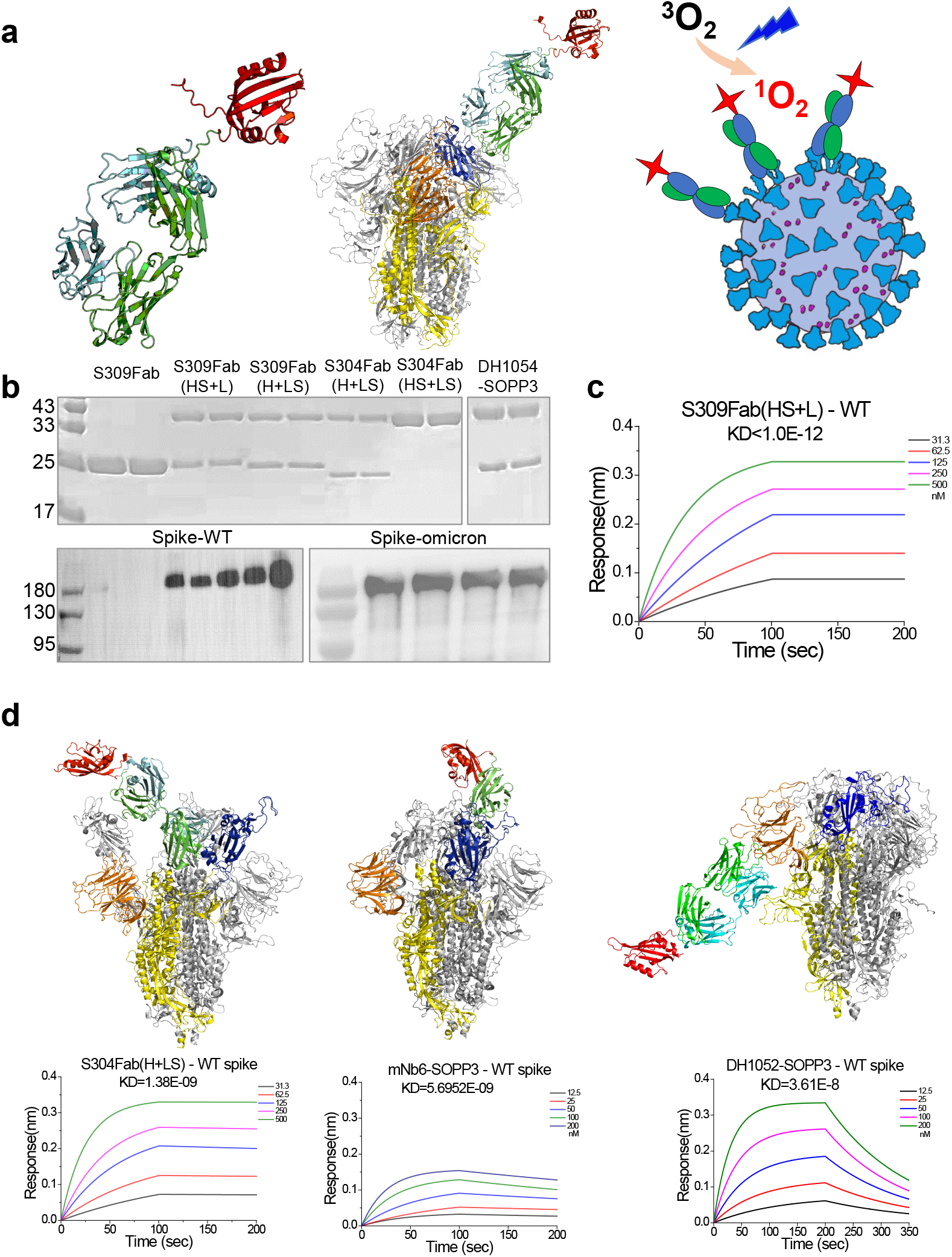
Construction, purification, and biochemical characterization of chimeric proteins containing a genetically encoded photosensitizer and spike-targeting antibodies. (**a**) Left, the S309Fab(HS+L) structure modeled by AlphaFold. Cyan, heavy chain; green, light chain; red, SOPP3. Middle, structure (model) of S309Fab(HS+L) in contact with the WT spike protein. Blue, RBD; orange, NTD. Right, schematic drawing showing the design of the targeted photodynamic neutralization of SARS-CoV-2. (**b**) Top, Coomassie blue staining of protein gels for S309Fab and Ab-SOPP3 fusion proteins. Bottom, Western blot analysis of purified WT and Omicron spike protein. (**c**) BioLayer Interferometry (BLI) assay of binding of S309Fab(HS+L) to WT spike. (**d**) Structures (model) of WT spike in complex with S304Fab(H+LS) (left), mNb6-SOPP3 (middle), or DH1052(H+LS) (right).

We first engineered a genetically encoded photosensitizer, SOPP3, to either the N- or C-terminus of each antibody (**Fig. 1b**). SOPP3 was developed from miniSOG (mini singlet oxygen generator), a protein of plant origin and binds tightly to flavin mononucleotide (FMN).^34^ FMN can effectively produce ^1^O_2_ with a high quantum efficiency (Φ=0.65).^35, 36^ Five mutations, W81L/H85N/M89I/Y98A/Q103V, were introduced into SOPP3 to improve the diffusion of molecular oxygen through the protein and quench unnecessary electron transfer reactions.^37^ The quantum yield of singlet oxygen by SOPP3 is 0.61 (under 21% O2), a value very close to that of FMN. Ab-SOPP3 fusion proteins were expressed in bacteria or mammalian cells and purified by chromatographic methods.

First, we used BioLayer Interferometry (BLI) to examine the binding between Ab-SOPP3 fusion proteins and the spike extracellular domain (ECD). S309Fab(HS+L), in which SOPP3 is attached to the C-terminus of the heavy chain, binds to the WT spike tightly, with the K_D_ value less than 1×10^-12^ μM (**Fig. 1c**). A similar strategy was adopted to construct and characterize other Ab-SOPP3 fusion proteins (**Fig. 1d;**). To examine the functionality of SOPP3 in the Ab-SOPP3 fusion protein, we checked photosensitized release of ^1^O_2_ using spectroscopic detection of the NIR luminescence signal centered at 1270 nm - a golden standard for the detection of ^1^O_2_. Rose Bengal, a popularly used photosensitizer for photodynamic generation of ^1^O_2_, was used as the positive control (**Fig. 2a;**). Indeed, robust 1270 nm signals could be detected from the SOPP3-mNb6 sample upon light excitation.

**Fig. 2.**
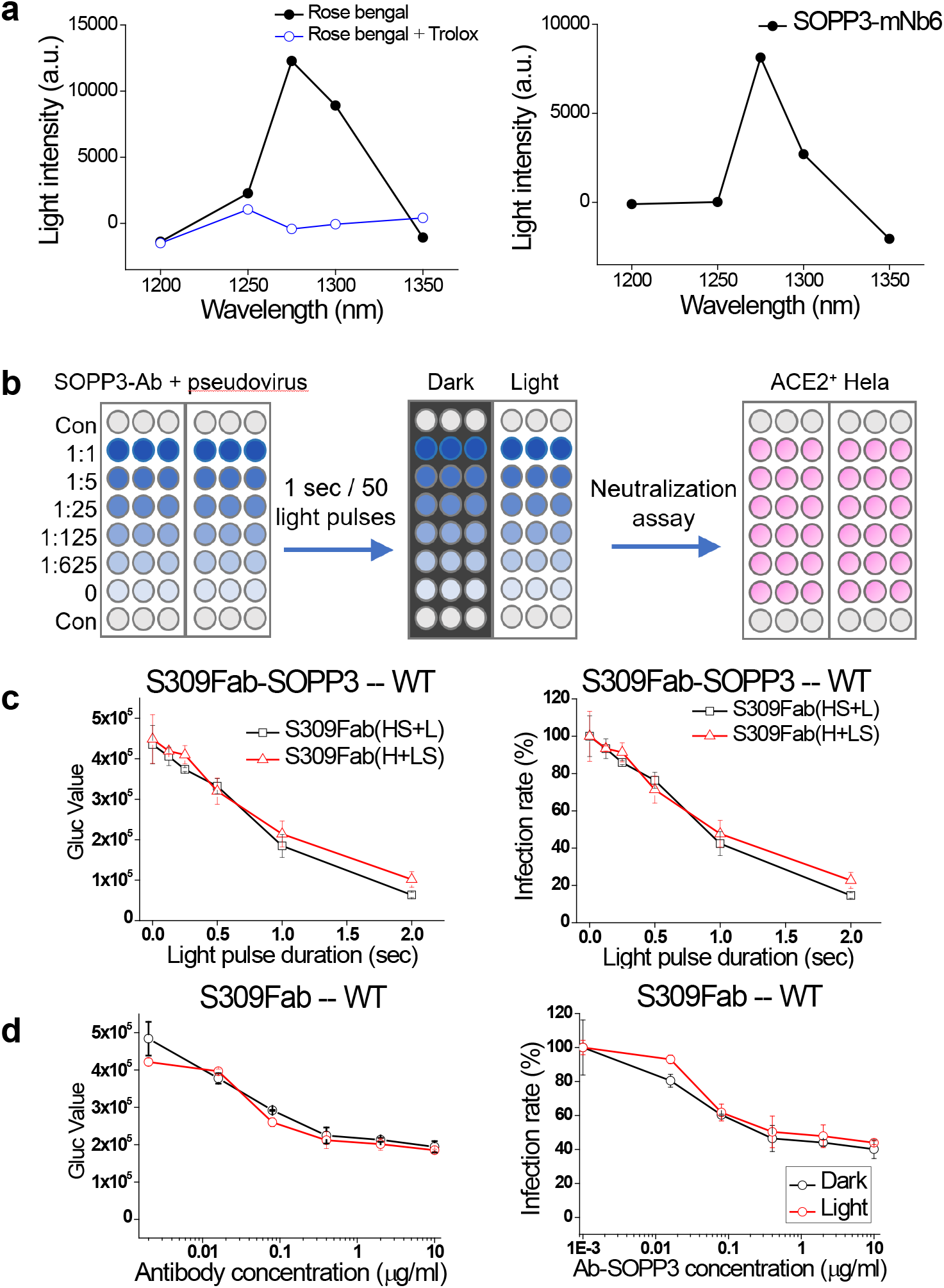
Near-infrared (NIR) spectroscopic detection of singlet oxygen (1O2) and setup of photodynamic neutralization assay. (**a**) Left, NIR signal of 10 μM rose bengal as the positive control (black). 5 mM Trolox (water-soluble vitamin E) added to quench ^1^O_2_ (blue). Right, NIR signal of SOPP3-mNb6. (**b**) Schematic drawing showing the setup of photodynamic neutralization assay. Serial dilutions of Ab-SOPP3 were incubated with SARS-CoV-2 at 37°C for 30 mins, followed by exposure to 50 light pulses or in the dark, and then were added to ACE2-expressing Hela cells. (**c**) Pseudovirion infection rate as a function of light pulse duration. Secretion of Gaussia luciferase in the medium was quantified by luminometry. Left, raw data without normalization. Right, all results were normalized to the value in the dark. Two different S309Fab-SOPP3 fusion constructs at the concentration of 500 ng/ml were tested. (**d**) S309Fab without SOPP3 attachment is not sensitive to light treatment. Left, raw data without normalization. Right, normalized results.

Next, we proceed to examine these SOPP3-tagged antibodies in the photodynamic neutralization of SARS-CoV-2 pseudovirions (**Fig. 2b**). Briefly, virion samples were mixed with serially diluted Ab-SOPP3 and then exposed to blue light pulses. As the negative control, a separate batch of Ab-SOPP3 and virion mixture was kept in the dark for the same duration of time. Then the samples were added to Hela cells overexpressing human ACE2. After 48 hours of incubation, the cell culture supernatant was collected for luminometry detection of luciferase level.

We started by optimizing the duration of light pulses applied to the mixture of Ab-SOPP3 and WT virions (**Fig. 2c**). Four Ab-SOPP3 proteins at the concentration of 500 ng/ml were used in the test: S309Fab(HS+L) (**Fig. 2d**), S309Fab(H+LS), SOPP3-mNb6, and mNb6-SOPP3 (similar results; data not shown). The mixture of virion and Ab-SOPP3 was exposed to 50 light pulses of different durations, from 0, 0.5, 1.0, 1.5, to 2.0 seconds. To approach effective photodynamic neutralization with minimal non-specific photodamages, we chose the duration of 1 second. Using S309Fab without the SOPP3 tag as the negative control, we confirmed 50 1-second light pulses had no observable effects on the viral infectibility (**Fig. 2d**). It is noteworthy that at concentrations above 1 μg/ml, the infection rate plateaued around 40%, which could be attributed to the absence of the Fc fragment (**Fig. 3a**).^30^

**Fig. 3.**
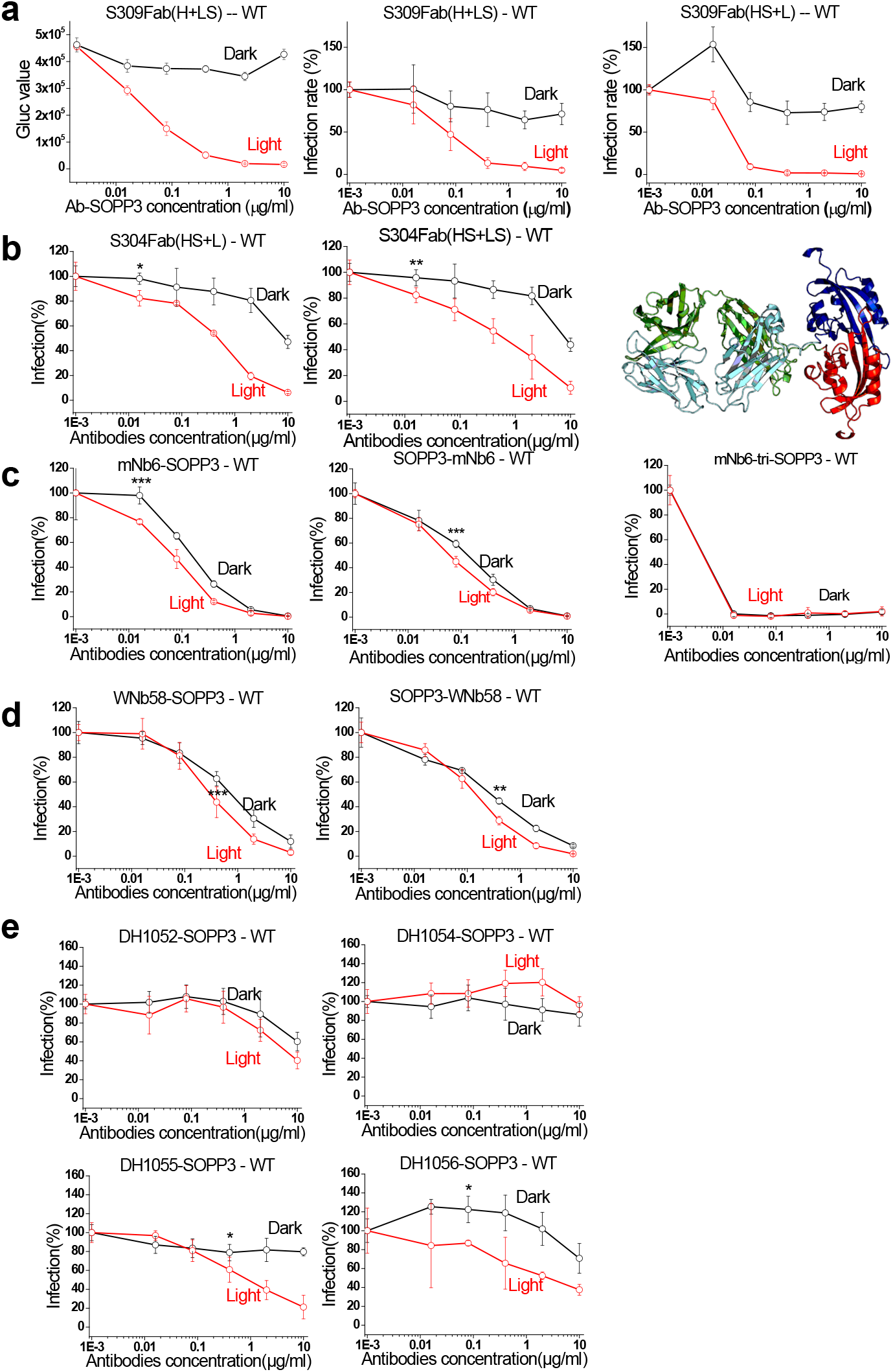
Photodynamic neutralization of the WT SARS-CoV-2 strain. (**a**) Left, raw data of S309Fab(H+LS). Middle, normalized results of S309Fab(H+LS). Right, normalized results of S309Fab(HS+L). Notice that at the concentration of 16 ng/ml and in the dark, S309Fab(HS+L) shows strong antibody-dependent enhancement (ADE) of infection. (**b**) Results of SOPP3 attached to S304. Left, S304Fab(HS+L). Middle, S304Fab(H+LS). Right, structure (model) of S304(HS+LS). (**c**) Results of SOPP3 attached to nanobodies against WT RBD. Left, SOPP3-mNb6, SOPP3 attached to the N-terminus of mNb6. Middle, mNb6-SOPP3, SOPP3 attached to the C-terminus of mNb6. Left, mNb6-tri-SOPP3, SOPP3 attached to the C-terminus of trimeric mNb6. Notice that mNb6-tri-SOPP3 is ultra-potent against WT virion. (**d**) Results of SOPP3 attached to non-neutralizing antibody WNb58. (**e**) Results of SOPP3 attached to NTD-targeting antibodies.

### Photodynamic neutralization of WT SARS-CoV-2

We then studied S309Fab(H+LS), in which SOPP3 is attached to the C-terminus of the light chain, in photodynamic neutralization of the WT strain. Compared to the original S309Fab fragment, S309Fab(H+LS) showed a weaker neutralization efficacy in the absence of light treatment (**Fig. 3a**, left, black trace). However, the application of light pulses significantly increased the neutralization efficacy, with the infection rate decreasing to close to zero at the concentration of 0.4 μg/ml (IC50: 68 ng/ml; **Fig. 3a**, left, red trace). Normalized results are shown in **Fig. 3a**, middle. The effects of photodynamic neutralization were more pronounced for S309Fab(HS+L), in which SOPP3 was attached to the heavy chain. At the concentration of 80 ng/ml, light pulses decreased the infection rate from 85.5 ± 11.2 % (N=6) to 9.2 ± 3.1 % (N=6) (**Fig. 3a**, right). Notably, at the concentration of 16 ng/ml, S309Fab(HS+L) produced a significant antibody-dependent enhancement (ADE) of infection, reflected in the infection rate increase to 153.8 ± 20.6 %.

According to the original report, the neutralization efficacy of S304 is marginal compared to S309 ^30^. Impressively, in our photodynamic neutralization assay, S304Fab-SOPP3 fusion proteins showed a significantly improved neutralization efficacy. We examined two constructs: S304Fab(HS+L) and S304Fab(HS+LS). Application of light pulses significantly improved the neutralization efficacy for both constructs. The IC50 values were decreased to a level around 50 ng/ml, with the infection rate reduced to close to zero at the concentration of 10 μg/ml (**Fig. 3b**).

For nanobodies, we examined six constructs: mNb6-SOPP3, SOPP3-mNb6, mNb6-tri-SOPP3, SOPP3-mNb6-tri, WNb58-SOPP3, and SOPP3-WNb58 (**Fig. 3c**). In the absence of light pulses, all six constructs effectively reduced the infection rate to close to zero. Application of light pulses slightly improved the neutralization efficacy of these Ab-nanobody fusion proteins. Notably, WNb58 is a non-neutralizing antibody according to the original report ^33^. However, we discovered that attachment of SOPP3 increased the neutralization efficacy of WNb58 (**Fig. 3d**).

To further explore the potential of this approach, we examined antibodies targeting the NTD domain of the WT spike protein, including DH1052, DH1054, DH1055, and DH1056, which according to the original report are all infection-enhancing antibodies.^32^ In the absence of light treatment, we didn’t observe any significant ADE in SOPP3-attached DH1052, DH1054, and DH1055 (**Fig. 3e**). For DH1052-SOPP3 and DH1054-SOPP3, application of light pulses produced marginal effects on the infection rate. The decreases in infection rate became more prominent for DH1055-SOPP3 and DH1056-SOPP3, especially at concentrations above 80 ng/ml. Notably, consistent with the original report of DH1056, DH1056-SOPP3 showed ADE at concentrations between 16 and 400 ng/ml.

### Photodynamic neutralization of the Delta variant

The Delta variant (B.1.617.2) examined in our system contains the following mutations: T19R, G142D, Δ156-157, R158G, L452R, T478K, D614G, P681R, D950N. In contrast to the observations with the WT strain, application of light pulses produced significant increases in the neutralization efficacy against the Delta strain for most Ab-SOPP3 fusion proteins, except mNb6-tri-S and S-mNb6-tri, two trimeric nanobodies showing high potency against Delta even in the dark.

Without the SOPP3 attachment and regardless of application of light pulses, S309Fab decreased the infection rate of the delta strain to a level around 40% (10 μg/ml), which is comparable to the effect on the WT strain. For S309Fab(H+LS) and S309Fab(HS+L), two fusion proteins containing SOPP3, application of light pulse significantly improved the efficacy of neutralization, with the IC50 decreased from 61 to 15 ng/ml and the infection rate reduced to almost zero at the concentration of 400 ng/ml (**Fig. 4a**).

**Fig. 4.**
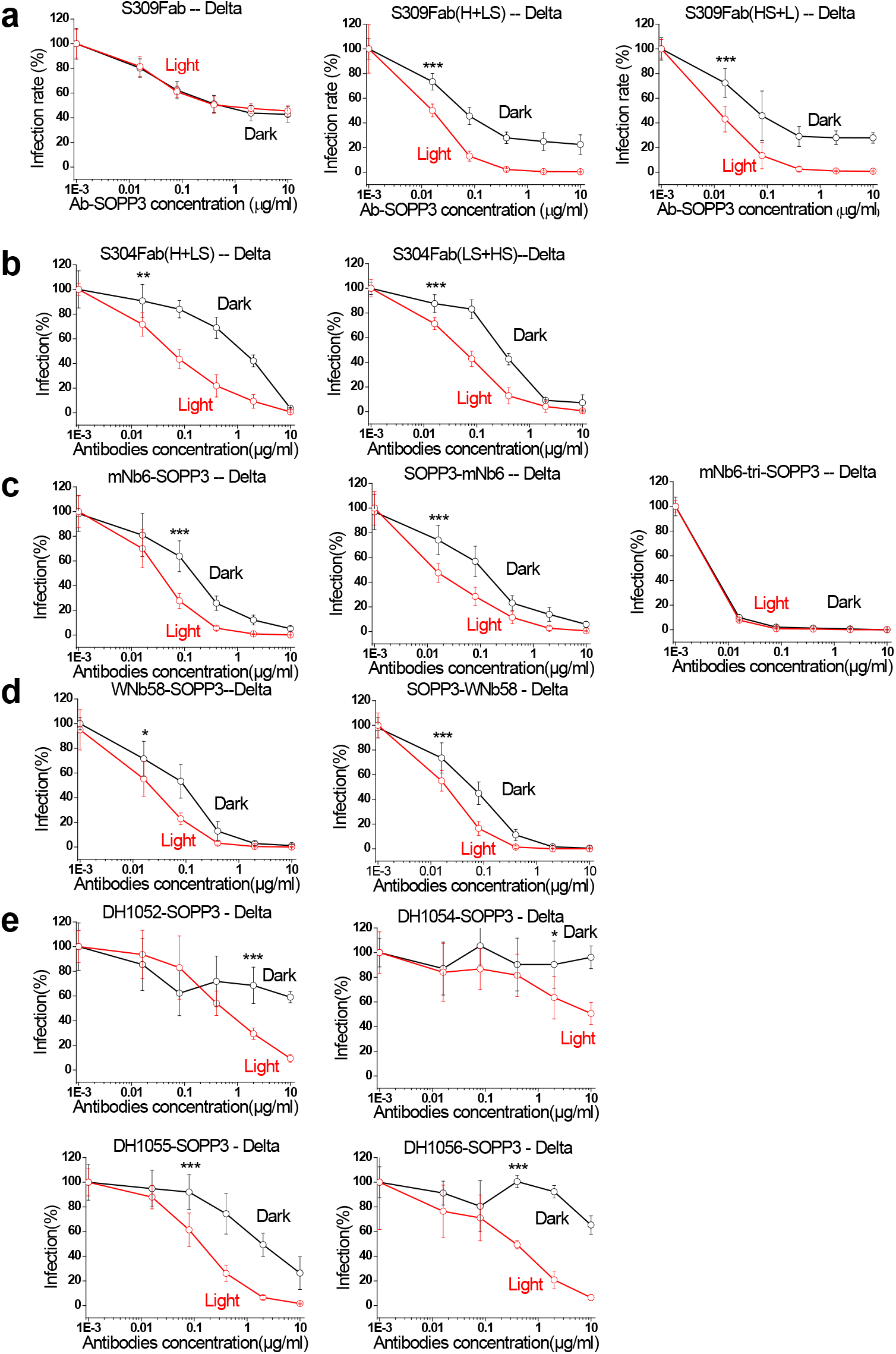
Photodynamic neutralization of the SARS-Cov-2 Delta variant. (**a**) Left, S309Fab. Light treatment had no effects on the neutralization efficacy of S309Fab against the Delta virion. Middle, S309Fab(H+LS). Right, S309Fab(HS+L). (**b**) Left, S304Fab(H+LS). Right, S304Fab(HS+LS). (**c**) Left, mNb6-SOPP3. Middle, SOPP3-mNb6. Right, mNb6-tri-SOPP3. (**d**) Left, WNb58-SOPP3. Right, SOPP3-WNb58. (**e**) Results of SOPP3-tagged antibodies targeting the NTD domain.

In the absence of light pulses, S304Fab(H+LS) and S304Fab(HS+L) could decrease the infection rate of the Delta strain to close to zero (10 μg/ml; **Fig. 4b**, black traces). Still, applying light pulses shifted the dose-response curve of both Ab-SOPP3 fusion proteins leftward and decreased the value of IC50 by more than 20 folds (**Fig. 4b**, red traces). Similar observations were obtained for mNb6-SOPP3 and SOPP3-mNb6, two nanobody constructs (**Fig. 4c**), WNb58-SOPP3 and SOPP3-WNb58, two RBD-targeting human antibody constructs (**Fig. 4d**), and four NTD targeting antibodies. Impressively, for DH1056-SOPP3, the infection rate of the Delta strain was decreased from 92.3 ± 5.1 % (N=6; dark) to 20.8 ± 7.2 % at the concentration of 2 μg/ml (N=6; after light pulses) (**Fig. 4e**). Taken together, these results highlight the effectiveness of this targeted photodynamic approach against the Delta SARS-CoV-2 strain.

### Photodynamic neutralization of the Omicron variant

According to recent publications, the Omicron variant carries extensive mutations in the spike protein and escapes most therapeutic antibodies, except VIR-7831 (Sotrovimab), which was developed from S309. We first applied BLI assay to examine the binding between Omicron spike and Ab-SOPP3 fusion proteins. As expected, we observed weakened binding to the omicron spike by S309, SOPP3-tagged S309, and SOPP3-tagged S304 (**Fig. 5a**). Then we examined the sensitivity of the Omicron virion to the procedure of photodynamic neutralization. For SOPP3-tagged S309Fab proteins, we observed a significant improvement in neutralization efficacy upon light treatment, reflected in the decrease in the infection rate when the concentration of S309Fab(H+LS) or S309Fab(HS+L) was above 80 ng/ml (**Fig. 5b**). For S304Fab(H+LS), application of light pulses elicited significant decreases in the infection rate at all concentrations tested (**Fig. 5b**). We obtained similar results from a separate batch of experiments. Since the experimental structures of the Omicron spike and the complexes of S309 and WT RBD have been published, we modeled the complex of Omicron and S309Fab(HS+L) (**Fig. 5c**). It appears that D336 (a point mutation unique to Omicron) and E337 in Omicron RBD contribute to the interaction with S309Fab. For other Ab-SOPP3 fusion proteins, BLI binding assays revealed no apparent binding to Omicron spike, and correspondingly, their neutralization against Omicron was completely diminished (**Fig. 5d, 5e**).

**Fig. 5.**
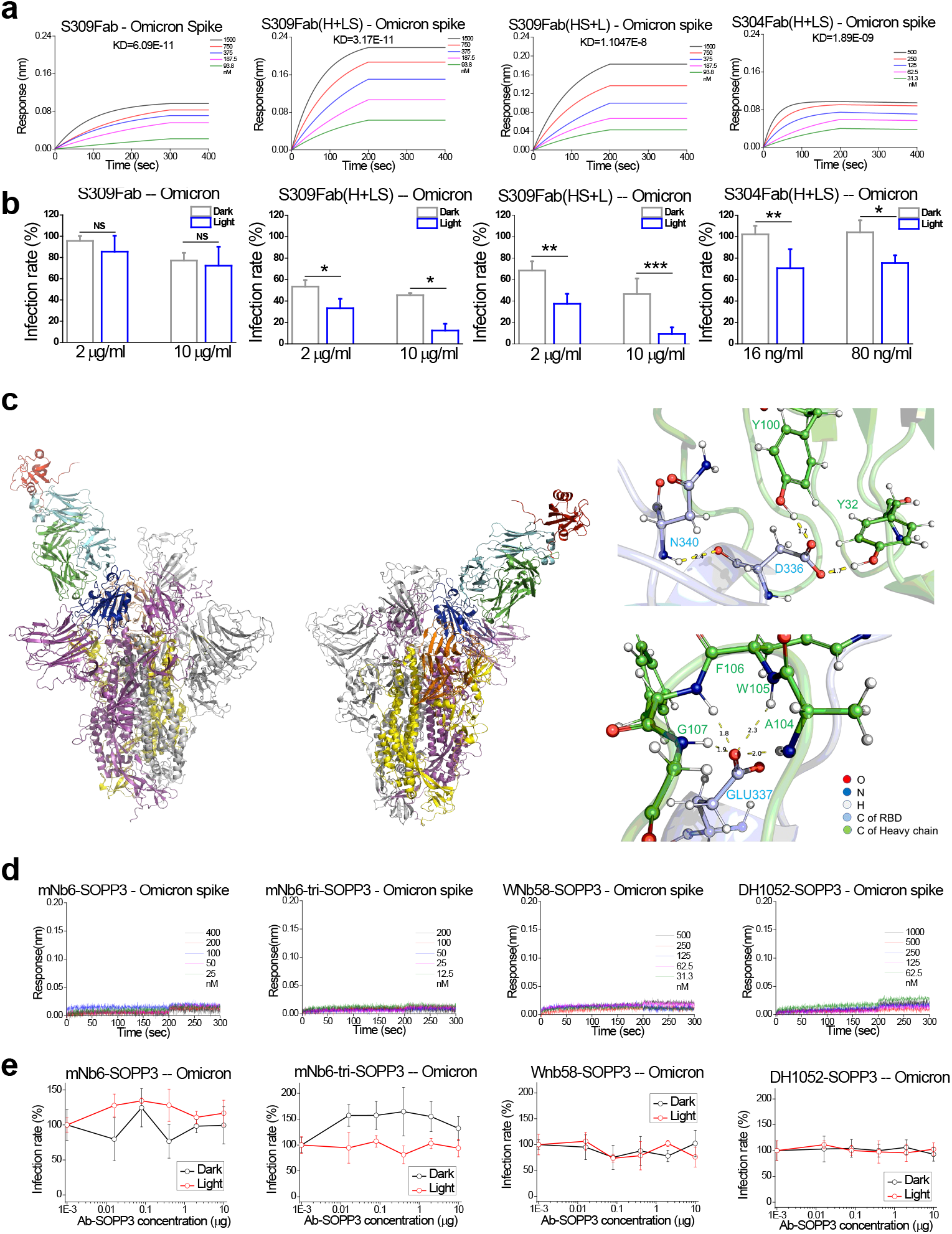
Photodynamic neutralization of the SARS-Cov-2 Omicron variant. (**a**) BLI assay of Ab-SOPP3 fusion proteins binding to the Omicron spike protein. From left to right, S309Fab, S309Fab(H+LS), S309(HS+L), S304(H+LS). (**b**) The corresponding neutralization results. From left to right, S309Fab, S309Fab(H+LS), S309(HS+L), S304(H+LS). Results of two concentrations of Ab-SOPP3 are shown. (**c**) Snapshot of a 10-ns MD simulation of Omicron spike in complex with S309Fab(HS+L). An extensive network of hydrogen bonds supports the interaction between Omicron RBD and S309. (**d**) BLI assay of Ab-SOPP3 fusion proteins binding to the Omicron spike protein. From left to right, mNb6-SOPP3, mNb6-tri-SOPP3, WNb58-SOPP3, DH1052(H+LS). None of these Ab-SOPP3 fusion proteins show obvious binding to the Omicron spike protein. (**e**) The corresponding neutralization results. From left to right, mNb6-SOPP3, mNb6-tri-SOPP3, WNb58-SOPP3, DH1052(H+LS). Consistent with the BLI binding assay, no obvious neutralization of the Omicron virion was observed for these fusion proteins.

### Results of mass spectrometric analysis

To explore the molecular nature of photodynamic modifications made to the spike protein, we applied mass spectrometry and analyzed the WT spike protein in complex with S309Fab(HS+L) (**Fig. 6a**). Identical preparations without light exposure were used as the negative control. We focused on residues in the spike protein showing a mass shift of +16 or +32 Da, corresponding to singly and doubly oxidizations, respectively (**Fig. 6b**). To validate the results, we sequentially repeated this experiment four times, all with separate and parallel negative controls. Interestingly, all modified residues are mainly located either in the N terminus of the spike protein, namely the NTD domain (Y144, Y145, H146, H207, H245, Y265) and the RBD domain (Y449, Y453, H519), and in the C-terminal trunk region (M1050, H1058, H1064). Among them, H146 and Y145 are two residues that stand out from four sets of experiment results and appear to form an “oxidation hotspot”. Interestingly, these two residues are located in the center of the antigenic “supersite” in the NTD domain recognized by many Nabs.^38, 39^ The NTD supersite refers to an epitope comprising residues from three loops: N1 (Q14–P26), N3 (L141–E156), and N5 (R246–A260) (**Fig. 6c**). Therefore, it is possible that modifications made to this antigenic supersite in NTD during the photodynamic process, in conjunction with modifications made to RBD and other regions in the spike protein, compromise the docking of the spike to ACE2 and consequently decrease the virulence of SAR-CoV-2.

**Fig. 6.**
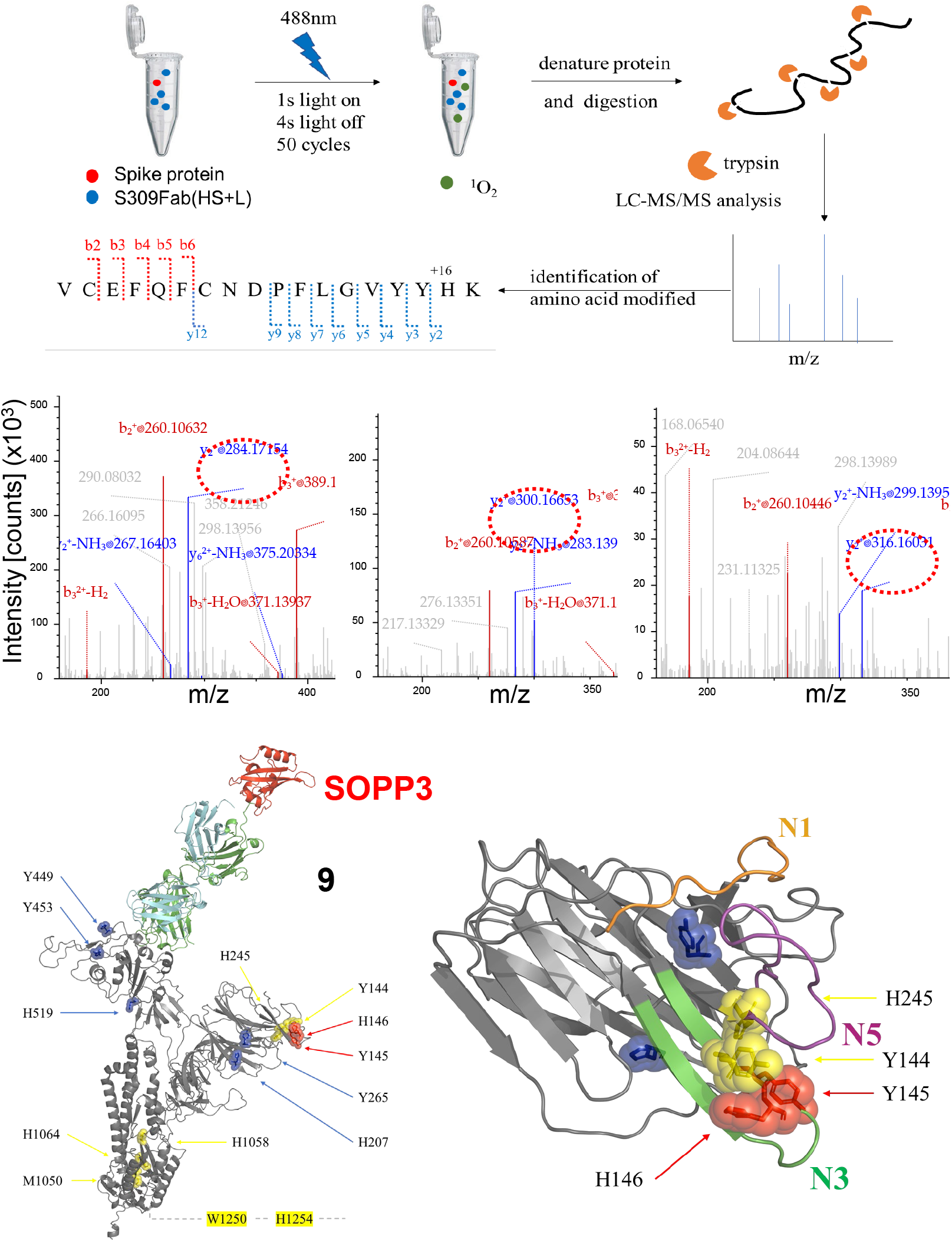
Mass-spec investigation of the sites and nature of oxidization in the spike protein elicited by the photodynamic process involving S309Fab(HS+L). (**a**) A schematic drawing showing the experimental procedure. (**b**) Mass spectrometry results for the peptide containing Y145 and H146. Red circles indicate modifications made to H146. From left to right, no light exposure, single oxygen addition (with light), and double oxygen addition (with light). (**c**) Left, structure (model) a WT spike protomer in complex with S309Fab(HS+L). The residues being identified for 2 (blue), 3 (yellow), or 4 (red) times out of a total of four experiments are shown. Right, a zoomed view over Y145 and H146, two residues located in the center of the “supersite” in NTD and oxidized during the photodynamic process. A supplementary video provides a rotating view over one S309Fab(HS+L) molecule in complex the trimeric WT spike protein.

To further explore the impact of photosensitizer position on the modification of the spike protein, we chose DH1052(H+LS), an Ab-SOPP3 fusion recognizing the NTD domain but did not show much neutralizing activity in pseudovirus neutralization assay. We repeated the experiments (complex formation; light treatment; mass spectroscopy) 7 times and summarized the results in Fig. S11 and S12. Comparing the results of S309(HS+L), a slightly different pattern of oxidization imprints was left on the spike in complex with DH1052(H+LS). As our understanding of ^1^O_2_ as a potentially effective signaling factor and how ^1^O_2_ reacts with amino acids within its reach are not clear, our results provide a vivid view at the molecular level regarding the photodynamic modification of protein molecules mediated by ^1^O_2_.

## DISCUSSION

Here we developed a targeted photodynamic approach to neutralize not only the WT SARS-CoV-2 strain but also the later emerged Delta and Omicron variants. Attaching a genetically encoded photosensitizer, SOPP3, effectively improved the neutralization efficacy of a list of antibodies of diverse sources and functions against the WT and the Delta strains. Non-NAbs are effective candidates for this strategy, especially the antibodies that bind to conserved regions outside RBD and thus do not produce any evolution pressure for SARS-CoV-2. For the Omicron strain, given its extensive mutations in spike, it was not surprising that significant binding escape was observed for most antibody-SOPP3 fusion proteins examined in our study. However, corresponding to the residual binding shown by S309 and S304 fusion proteins, application of light pulses still improved the efficacy of both Ab-SOPP3 constructs against the Omicron strain.

One primary goal of the current study is to identify non-NAbs that target highly conserved regions in the spike protein and then utilize them in the photodynamic neutralization against SARS-CoV-2 and related viruses. This strategy can potentially expand the applicability of hundreds of antibodies against the continuously emerging immune escape variants. Indeed, our results of non-neutralizing antibodies and the antibodies targeting the NTD domain, such as DH1055 and DH1056, against the Delta strain validated this strategy. However, extensive escape mutations residing in the Omicron spike obliterate the binding of most therapeutic antibodies currently available. The moderate effects observed with S309Fab-SOPP3 and S304Fab(H+LS) against the Omicron strain necessitate the continuous efforts to identify broadly recognizing but not necessarily neutralizing antibodies. Non-NAbs do not elicit escape pressure on the virus, and among them, those target conserved regions in spike and thus have a broad recognition spectrum will be very useful for this targeted photodynamic approach.

Singlet oxygen is believed to play a central In the photodynamic elimination of malignant cells and pathogens.^40^ Our spectroscopic measurement of infrared luminescence signal centered at 1270 nm confirmed singlet oxygen generation by Ab-SOPP3 fusion proteins. Furthermore, to explore the chemical nature of modifications made to the spike protein, we used mass spectrometric analysis and identified targeted residues from light-exposed samples. The location of two significant residues (Y145 and H146 in NTD) is in line with the positioning of SOPP3 as S309Fab(HS+L) targets explicitly the nearby RBD domain. Moreover, our identification of a histidine residue as the most significant site is consistent with previous reports (mostly carried out in the gas phase) that histidine has the highest reaction rate to ^1^O_2_ (His, 5; Trp, 3; Met, 2; Cys, 0.9; Tyr, 0.8; in 10^7^ M^-1^s^-1^).^41, 42^ Conversely, our mass-spectrometric identification of H146 and neighboring Y144 and Y145 validates the significant role played by ^1^O_2_ in photodynamic processes.^43, 44^ Remarkably, H146 is highly conserved in the NTD of SARS-CoV-2 variants, across the WT, Delta, and particularly the immune escape Omicron strains. The spike protein is intrinsically allosteric and therefore modifications to the region outside RBD can remotely affect the binding of spike to its receptor on host cells like ACE2.^45, 46^ The perfect overlap between this “oxidation hotspot” and the antigenic “supersite” in the NTD ought to underlie the effectiveness of this photodynamic approach against SARS-CoV-2.

Vaccines in combination with neutralizing antibodies against SARS-CoV-2 carry great hope in the elimination of COVID-19. However, the relentless emergence of variants of concern presses for novel countermeasures. We developed a targeted photodynamic approach to neutralize SARS-CoV-2 by engineering a genetically encoded photosensitizer to a diverse list of antibodies targeting the WT spike protein. Pseudovirus neutralization assay confirmed the effectiveness of this photodynamic approach against not only the WT but also the Delta and the immune escape Omicron strains. Mass spectrometric analysis pinpointed the oxidation sites within the spike protein. Our study broadened the application potential of the antibodies against SAR-CoV-2 and paves the way for targeted photodynamic therapy in the treatment of infectious diseases. We anticipate the major technical hurdle for this approach to be used in clinical pratic is the delivery of light to deeper tissue, including the lung, in human body. Photosensitizers in the red and infra-red range or even sonosensitizers would form the possible solution.

## MATERIALS AND METHODS

### Molecular cloning, expression, and purification of recombinant proteins

The cDNAs of SOPP3 and antibodies were codon-optimized and commercially synthesized. Fusion PCR reactions were used to link two cDNAs together, with a linker sequence added in the frame between DNA fragments. The PCR products were cloned into the expression vector by standard recombination or restriction enzyme cut–ligation method.

The cDNA fragment encoding nanobody-SOPP3 fusion protein was cloned into the pSMT3 expression vector and transformed into *E. Coli* BL21 cells. Unnecessary light exposure during protein expression and purification was reduced to a minimum. When the bacteria growth reached the exponential phase, IPTG (0.5~1 mM) was added to the cell culture at 30 *°C* (or 18 °C) to induce protein expression. Cells were grown at 30 *°C* for 3 hours (or 18 *°C* for 12 h) and harvested by centrifugation (6000 rpm, 4°C, 8 minutes). Cell pellets were resuspended in lysis buffer (300 mM NaCl) and lysed by sonication on ice (2s on 3s off, 4min). The soluble fraction was separated from the insoluble components by centrifugation (15000 ×g, 4 °C, 1h) and then loaded onto the His-Trap column (5 ml). After protein elution by imidazole with a gradient of 40 to 240 mM, the His-tag was removed during the following dialysis step. Protein samples were further purified by Ion Exchange Chromatography and Size-exclusion Chromatography. Final products were aliquoted, snap-frozen in liquid nitrogen, and stored at −80°C.

Human antibody-SOPP3 fusion proteins and the ECD of spike were expressed in suspension 293F cells. Briefly, the plasmids encoding recombinant proteins were transiently transfected into 293F cells at the density of 1.5 × 10^6^ cells/ml. Unnecessary light exposure during protein expression and purification was reduced to a minimum. Cells were maintained at 37°C, 8% CO2, with the rotation speed set as 170 rpm. A daily sampling of the cell culture was collected, and the expression of the target protein was evaluated by western blot analysis. After 48 hours of incubation, cells were collected by centrifugation at 1000 rpm for 15 minutes. while the spike-expressing cells were collected after 72h of the identical incubation.

### BioLayer interferometry measurement of antibody-SOPP3 binding to Spike ECD

An Octet RED384 system was used for measuring the binding between purified WT or Omicron spike ECD and antibody-SOPP3. The purified spike ECD (10 mg/ml) contains an SBP-tag in the C-terminus and was captured by a Streptavidin-specific biosensor (#18-5019). The loading, association, and dissociation buffer contains 150 mM NaCl, 10 mM HEPES, 0.1% Tween-20, 0.1% BSA (pH=8.0). The regeneration buffer contains 10 mM glycine (pH 2.0).

### Singlet oxygen NIR luminescence measurement

A schematic of the experimental setup used to measure ^1^O_2_ luminescence is shown in Fig. S5. A 10 mm pathlength quartz cuvette was used as the solution container (Allrenta, China). A laser source (450/505/532 nm; OX-45001X by OXLasers, China) was used to excite photosensitizers. An 1150 nm long-pass filter (Thorlabs, USA) was used to block out the fluorescence from the sample and the scattering excitation light. Spectral discrimination of the detected signal was achieved using a set of five narrow-band filters centered at 1200, 1250, 1275, 1300, and 1350 nm (OD4 blocking, 45-nm full-width at half-maximum or FWHM; Edmund, USA) and mounted on a 6-position motorized filter wheel (FW102C, Thorlabs, USA). The ^1^O_2_ luminescence signal was detected by a NIR-PMT (H12694A-45-C4, Hamamatsu, Japan). The operating voltage of the PMT was set to −700 or −750 V. The PMT output was amplified and converted to a voltage pulse by an electrophysiology amplifier (A-M 2400, A-M Systems, USA). The Clampfit program was used for offline analysis of the NIR signals. Antibody-SOPP3 samples were diluted in the buffer containing 25 mM HEPES, 200 mM NaCl, pH 7.5) to a concentration of 1.5 mM.

### Mass spectrometric (MS) analysis

#### Chemicals

Ammonium bicarbonate, TCEP (tris(2-carboxyethyl)phosphine), dithiothreitol (DTT), IAA (iodoacetamide), TFA (trifluoroacetic acid), FA (formic acid) and ACN (acetonitrile) were purchased from Sigma-Aldrich. Trypsin Gold (mass spectrometry grade) was purchased from Promega.

#### Protocol for reduction, alkylation, and trypsin digestion

Protein samples were dissolved in 50 mM ammonium bicarbonate and 2-4 M urea, reduced with 10 mM TCEP at 37°C for 60 min and alkylated with 20 mM IAA at RT in the dark for 30 min. Then, the protein sample was diluted and incubated trypsin at 37 °C for 16 h. A C18 spin column (Cat. 89870, Pierce, USA) was used for desalting. Samples were dried and then dissolved in 0.1% FA. After centrifugation at 14000 g for 5 min, the supernatant was collected for MS analysis.

#### Nano-liquid chromatography and MS

The desalted protein digest was fractionated by nano-liquid chromatography–MS (nano-LC–MS) using an EASY-nLC 1000 system (Thermo Scientific). A C18 Acclaim PepMap RSLC analytical column (75 μm × 250 mm, 2 μm, 100Å) with a C18 nano Viper trap-column (0.3 mm ×5 mm, Thermo Fisher Scientific) was used for peptide elution and separation, with the flow rate set at 300 nl/min. The mobile phase contained buffer A (0.1% formic acid) and B (0.1% formic acid, 80% acetonitrile). The gradient was set as following: 0 min: 5% B; 90 min, 25% B; 105 min, 40% B; 110 min, 90% B; 117 min, 90% B; 120 min, 5%. MS data were then acquired with a Q Exactive™ Hybrid Quadrupole-Orbitrap™ Mass Spectrometer (Thermo Scientific) in positive mode, with the following settings: 1) MS1 scan range 400 and 1,600 m/z, resolution at 70,000, automatic gain control (AGC) 1e6 and maximum injection time 50 msec; 2) The collision energy was set at 32% and orbitrap was used for MS2 scan as well; 3) MS2 scan range was auto defined with resolution at 17,500, isolation window 1.8 m/z, AGC 5e4 and maximum injection time 100 ms; 4) Exclusion window was set for 25 sec; 3) The intensity threshold was set at 8e3.

Data analysis was carried out with Proteome Discoverer 2.5 using standard settings for each instrument type and searched against a *Homo sapiens* database downloaded from UniProt in 2021. A peptide tolerance of 10 ppm and fragment ion tolerance of 20 ppm was used. Carbamidomethylation of cysteine was specified as fixed modification, while oxidation of methionine, tyrosine, histidine, tryptophan of protein N termini were set as variable modifications. The false discovery rate was set to 0.01 for proteins and peptide spectrum match.

### Production of pseudotyped viruses for neutralization analysis

SARS-CoV-2 pseudotyped viruses of the WT strain and Delta and Omicron variants were produced for neutralization assay.^47^ Briefly, 293T cells were seeded at 30% density in 150 mm dishes 12-15 hours before transfection. Transfection was performed when the cell density reached 75-80%. Cells in each dish were transfected with 67.5 μg of polyethyleneimine (PEI) Max 40K (#40816ES03, Yeasen Inc.) and a mixture of plasmids encoding the spike protein (3.15 μg), the murine leukemia virus (MLV) gag, and pol proteins (15.75 μg), and the luciferase reporter (15.75 μg). Eight hours after transfection, the cell culture medium was changed to fresh medium containing 2% FBS and 25 mM HEPES. The cell culture supernatant was collected at 40-48 hours and 60-72 hours after transfection, centrifuged at 2000×g for 10 min and filtered through a 0.45 μm filter to remove cell debris. SARS-CoV-2 pseudotyped virions were concentrated using a centrifugal filter unit with a 100 kDa cut-off (Amicon UItra-15) and stored at −80°C.

### SARS-CoV-2 neutralization activity assay

Hela-ACE2 cells were seeded in 96-well plates at the density of 2×10^3^ cells per well. The SARS-CoV-2 pseudovirus was mixed with serially diluted Ab-SOPP3 fusion proteins and incubated at 37 °C for 30-60 min. After the incubation, the mixture in the light group was placed under a light source for light pulse irradiation, and the negative control group was protected from light throughout the process. After the light treatment, the mixture was added to Hela cells overexpressing ACE2 plated in a 96-well plate at the density of about 80%. After 15-17 hours of incubation, the medium was replaced with DMEM with 2% FBS. Gluc detection was performed at 48 hr using the Luciferase Assay System (Promega, Cat. #E1500).

### Protein structure modeling

#### Ab-SOPP3 fusion protein

Structures of Ab-SOPP3 fusion protein were built by AlphaFold2 and the extension AlphaFold-Multimer.^48^ The code and parameters were from DeepMind and used in the computer modeling of structures by AlphaFold neural network. Flexible linkers such as “GSASG” were added between SOPP3 and antibodies to reduce spatial hindrance between them.

#### Spike – Ab-SOPP3 complexes

The Modeller program was used to model the Spike protein in complex with Ab-SOPP3 fusion protein.^49^ The following structures involving SARS-CoV-2 spike were used: 6WPS, Spike-S309Fab(H+LS); 7LAB, Spike-DH1052(H+LS); 7JW0, Spike-S304(H+LS); 7KKL, Spike-mNb6-SOPP3.

#### Molecular dynamics simulation of Omicron-Ab-SOPP3 complex

The GROMACS program (2021.5 package) was used in the MD simulation of Omicron spike (reference PDB ID: 7WK3) in complex with S309Fab(HS+L).^50^ The Amber ff14sb force field and TIP3P water were used. Standard MD parameters were used in the simulation. The minimum distance between the protein and the boundary of the simulation box was set as 15Å. NaCl at the concentration of 150 mM was added to the system. The final production run was preceded by three energy minimization steps, one NVT and one NPT position-restrained MD runs.

### Data Analysis

The experimental data in the text and figures are expressed as mean ± SD. Student t-test (non-paired) was used to compare two sets of infection rates (dark vs. light) at a certain concentration of SOPP-Ab. The p-value is plotted in the figure as: ns, p> 0.05; *, p ≤ 0.05; **, p ≤ 0.01; ***, p ≤ 0.001.

## ACKNOWLEDGMENTS

We acknowledge the technical support and contribution to this project at earlier stages from Jizhong Han, Junhui Lai, Xiaoxi Li, Yi Li, and Ruohan Zhang. L. Zhou is supported by fundings from the National Natural Science Foundation of China (#32171150) and Shenzhen Bay Laboratory. We thank Dr. Xiaoliang Sunney Xie from Peking University and Dr. Qiang Zhou from Westlake University for the cDNA clones of WT and Omicron spike proteins. We thank Dr. Guocai Zhong for the generous help with the neutralization assay and related DNA plasmids. We thank Dr. Haidi Yin from Shenzhen Bay Laboratory Mass Spectrometry Core Facility for the MS analysis.

## Author contributions

Yao, R: molecular cloning, protein purification, and biochemistry, pseudovirus neutralization assay, data analysis and manuscript writing; Hou, J: molecular cloning, protein purification and biochemistry, NIR detection, data analysis, and manuscript writing; Zhang, X: protein modeling, data analysis, and manuscript writing; Wu, Q: molecular cloning, protein purification and biochemistry; Liu, Q: study design, data analysis and manuscript writing; Zhou, L: study design, data analysis and manuscript writing.

## Conflict of interest

none

## Notes

### Competing Interest Statement

The authors have declared no competing interest.

